# Hunger, displacement or play? Object manipulation behaviour in Asian small-clawed otters (*Aonyx cinereus*)

**DOI:** 10.1101/2025.11.14.688474

**Authors:** Iwan S. Evans, Andrew S. Cooke, Jonathan Cooper, Beth A Ventura

**Affiliations:** School of Natural Sciences, University of Lincoln, Lincoln, UK; College of Veterinary Medicine, Michigan State University, East Lansing, MI, USA

**Keywords:** zoo animal behaviour, animal welfare, ethogram, stone handling, function, mustelid

## Abstract

Asian small-clawed otters (*Aonyx cinereus*) engage in intricate activities using their paws for purposes such as food extraction, exploration, and tactile communication within their social group. However, some expressions of these behaviours do not appear to be immediately functional in captive populations. Here, we explore the expression of object manipulation behaviours (OMBs), such as rock manipulation (or ‘juggling’); the factors that may be driving their expression; and what OMBs may represent. A troop of six otters was observed at Lincolnshire Wildlife Park (United Kingdom) using camera-traps placed at three key locations in their enclosure: outside of their den, at the main feeding area, and at a rock-garden foraging area. A total of 2895 video clips were analysed using a hierarchical ethogram consisting of the behavioural categories of locomotion, stationary, vocalisation, physical affiliative, sustenance, and OMBs. The most commonly observed behaviours were walking (21.5%), lying (20.8%) and feeding (12.2%). OMBs were observed in 3.9% of total observations. Otters performed OMBs significantly more often outside their den than at the other locations. Grooming was also more frequent at this location. There was a negative association between the expression of OMBs and air temperature. Results suggest that in this otter population, OMBs were more likely an expression of play behaviour, contrasting findings reported elsewhere that rock juggling may be driven by hunger.

## 1 Introduction

Asian small-clawed otters (*Aonyx cinereus*) are the smallest species of otter. They are native to Southern Asia but are found in zoological institutions worldwide. In the wild, they are extractive feeders, typically feeding on molluscs, crabs and other aquatic animals (Kruuk, 2006; Heap et al. 2008). They are adapted to be relatively dexterous and have been observed using their paws for activities such as extracting and handling food and non-food objects in captivity (Frick et al. 2016; Manns et al. 2018).

Object manipulation behaviours (OMBs) have been observed across a wide variety of species for the purposes of play, learning, feeding/forage, nest building, and courtship (Sugasawa et al. 2021). These purposes often overlap, for example, learning through play to develop foraging skills. Allison et al. (2020) explored ‘rock juggling’ in smooth-coated otters (*Lutrogale pespicillata*) and Asian small-clawed otters (*Aonyx cinereus*) and suggested that ‘rock juggling’ may be misdirected behaviour when hungry, as they observed these behaviours more commonly at times when they predicted otters to be hungry. They also noted no relationship between rock juggling and the development of foraging skills but did not rule out the potential for rock-juggling to be a form of play behaviour associated with behavioural development.

### 1.1 Objectives

The primary aim of the study was to gain insight into the function and drivers of object manipulation in Asian small-clawed otters. This was achieved by:

- Univariate analysis of the spatial distribution of OMBs.
- Multivariate analysis of the spatial distribution of categories of behaviours, allowing for associations between locations and behaviours to be explored.
- In this respect, we were particularly interested in the relationship between OMBs and feeding/foraging behaviour.

## 2 Methods

### 2.1 Study site

All study procedures were evaluated and approved by the University of Lincoln Research Ethics Committee (UoL2023_13874).

The study site was the Asian small-clawed otter enclosure at Lincolnshire Wildlife Park (LWP), UK (53.0818, 0.1454). LWP is set in approximately 10 ha of land and houses a variety of birds, mammals and reptiles. The otter enclosure (Figure 1) is located near the centre of LWP along the main visitor path, adjacent to slender meerkats (*Suricata suricatta*), lowland tapir (*Tapirus terrestris*) and Sulcata tortoises (*Centrochelys sulcata*). The enclosure is primarily outdoors with an area of approximately 646m^2^ and is comprised of predominantly grassy ground with a pool in the centre of the enclosure, with reeds towards the rear and open water at the front, and a wooden bridge transecting the water. Towards the front of the enclosure, where visitors typically view (Viewing Area 1 in Figure 1), there is a large log and a pebble/rock garden. This location is where the otters are normally fed, with food items either scatter fed, placed on the log, or hidden amongst the pebbles of the rock garden. To the right of the enclosure, there is a raised wooden den accessed by a short ramp mostly obscured from visitors, though there is a secondary, less frequently used viewing area near it (Viewing Area 2).

**Figure 1.**
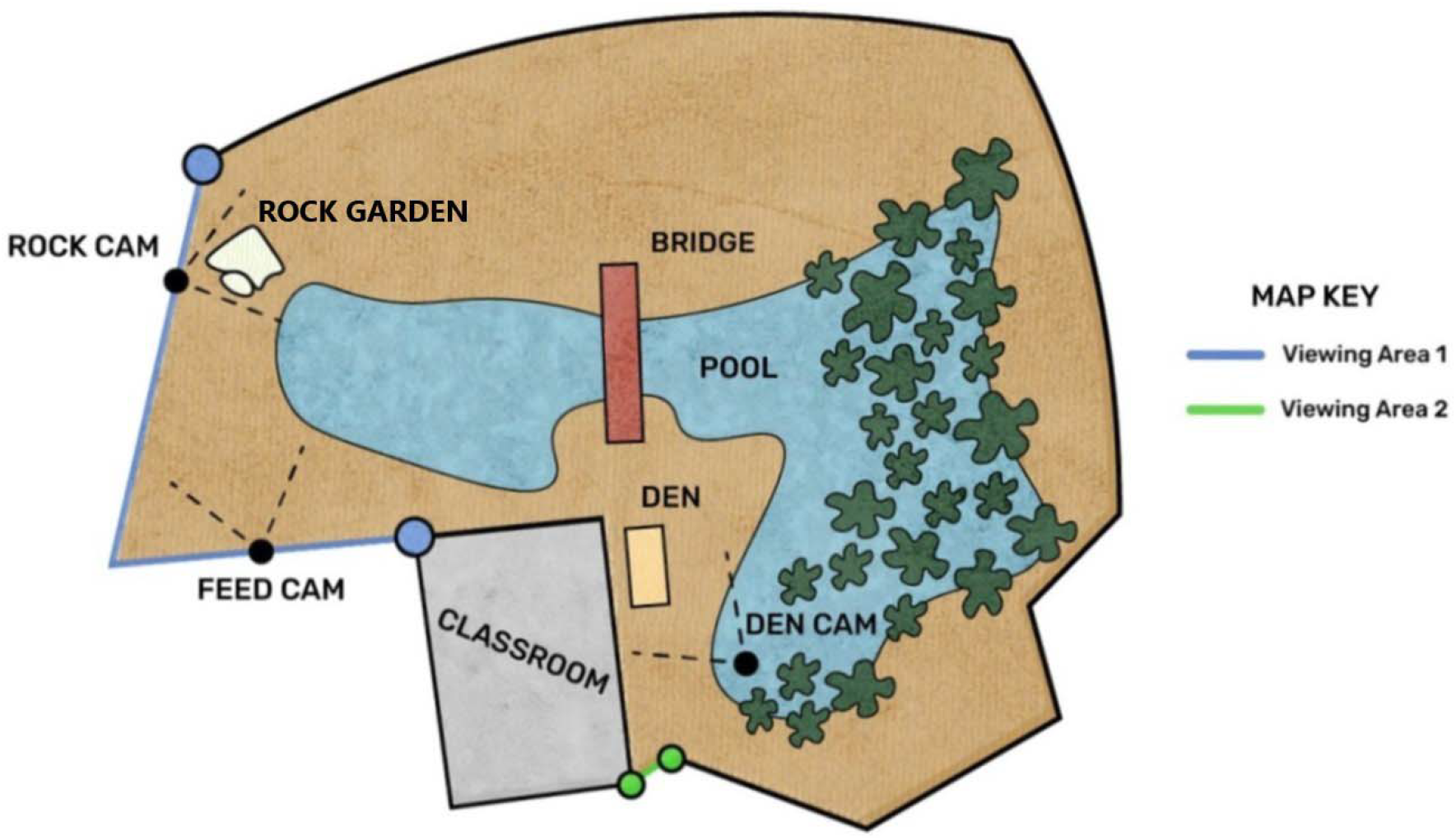
Diagram of the Asian small-clawed otter (646m^2^) enclosure with key features and camera locations annotated. Diagram is not to exact scale.

### 2.2 Study population

The population was comprised of six individuals: two mature adults (one male, one female, ∼8 yrs of age) and four young adults (three males and one female, ∼ 2 yrs) who are offspring of the mature female. The mature adults moved to LWP in 2020, having been relocated from Tamar Otter and Wildlife Park, UK (following the closure of their otter facilities). The young adults were all born at LWP in 2021.

Otters’ husbandry, and in particular their feeding regime, were consistent with guidelines set out by the IUCN/SSC Otter Specialist Group for Asian small-clawed otters (Heap et al. 2008). Otters were typically fed four times per day, usually between 0900 hrs-1000 hrs, 1100 - 1200 hrs, 1300 -1400 hrs, and 1600 – 1700 hrs. Their first morning meal normally consisted of eggs and (dead) chicks which were provided by hand feeding by a keeper. Remaining meals consisted of scatter feeding in the front enclosure area, which was the most visible area to visitors. Scatter feeding typically included beef or pork with a combination of mussels, some form of whole or chopped fish, and plant-based food such as celery or grapes. During scatter feeds some food items could be hidden around the enclosure, including the rock garden, to encourage searching or foraging behaviour.

### 2.3 Behaviour observations

#### 2.3.1 Video recording

Camera traps (Browning Recon Edge) were used to record behaviours for a total of 30 days across the periods 25/05/2023 to 30/05/2023 (5 days) and 17/07/2023 to 17/09/2023 (25 days). Cameras were motion-activated and recorded for 20s in 1080p with audio, with a 1s trigger interval. Cameras were placed approximately 30cm off the ground in three locations (Figure 1 for locations; Figure 2 for field of view):

1. **Rock Cam** was positioned to capture the area surrounding the rock garden at the front of the enclosure, where feeding typically occurred.
2. **Feed Cam** was positioned toward the area in which the otters were provided with their food near the front of the enclosure and captured a larger field of view.
3. **Den Cam** was placed near the den, away from the primary visitor viewing area.

**Figure 2.**
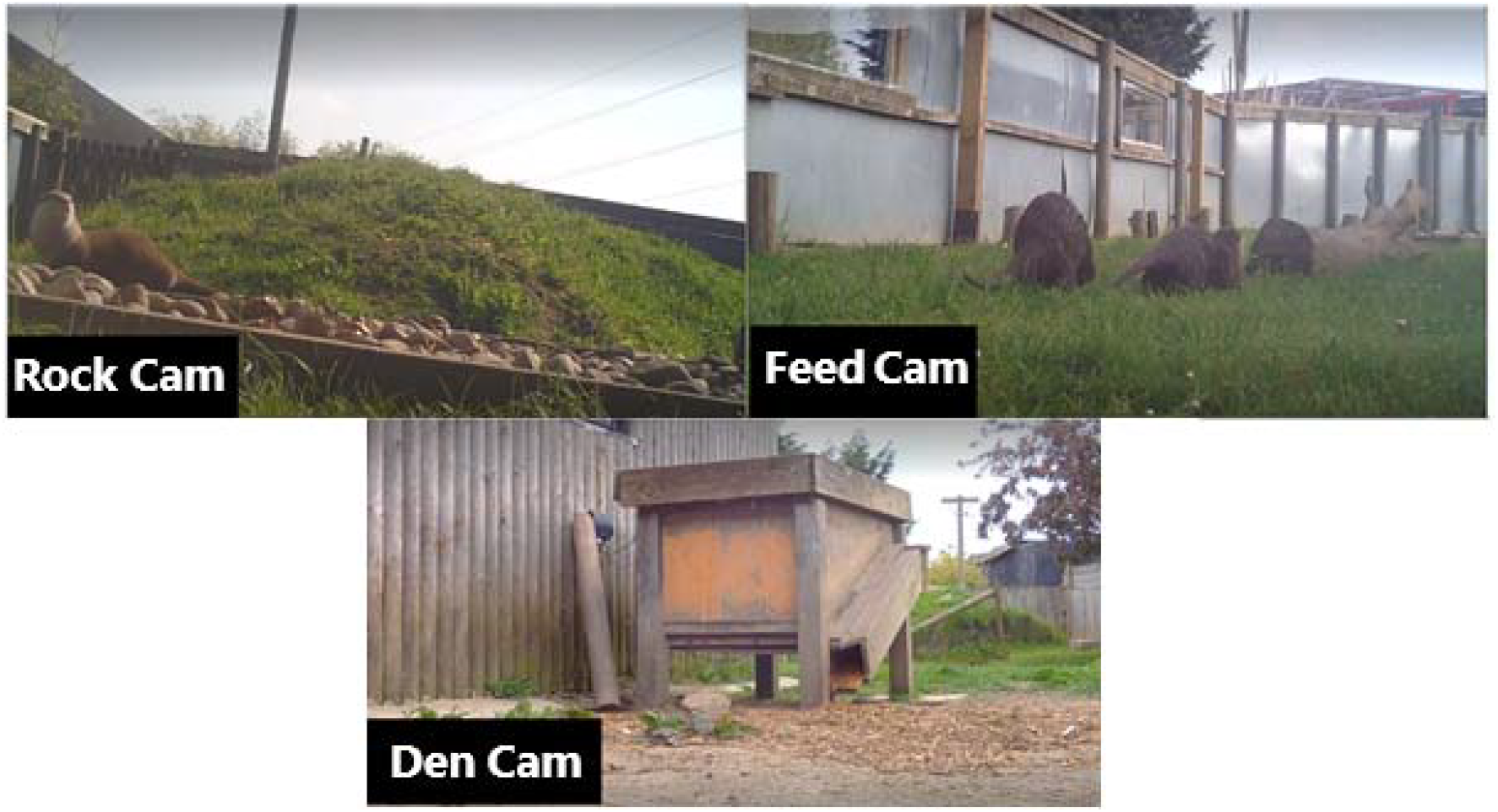
Video stills from the three camera traps set to capture behaviour in an Asian small-clawed otter zoo enclosure housing 6 otters.

These locations were specifically selected as key areas based on reports from LWP staff of where the otters were fed and where they had been seen to partake in object manipulation.

#### 2.3.2 Ethogram

An ethogram was developed and adapted from Allison et al. (2020) (Table 1). The primary modifications to that ethogram were the reclassification of handling behaviours to ‘object manipulation behaviours’ (OMBs). This is a broader umbrella category that does not focus specifically on rocks or ‘juggling’. It includes a variety of rock and straw manipulation behaviours not formally defined elsewhere. Video examples of OMBs from this study are available online (https://youtu.be/fITCIzKb_JY).

**Table 1.**
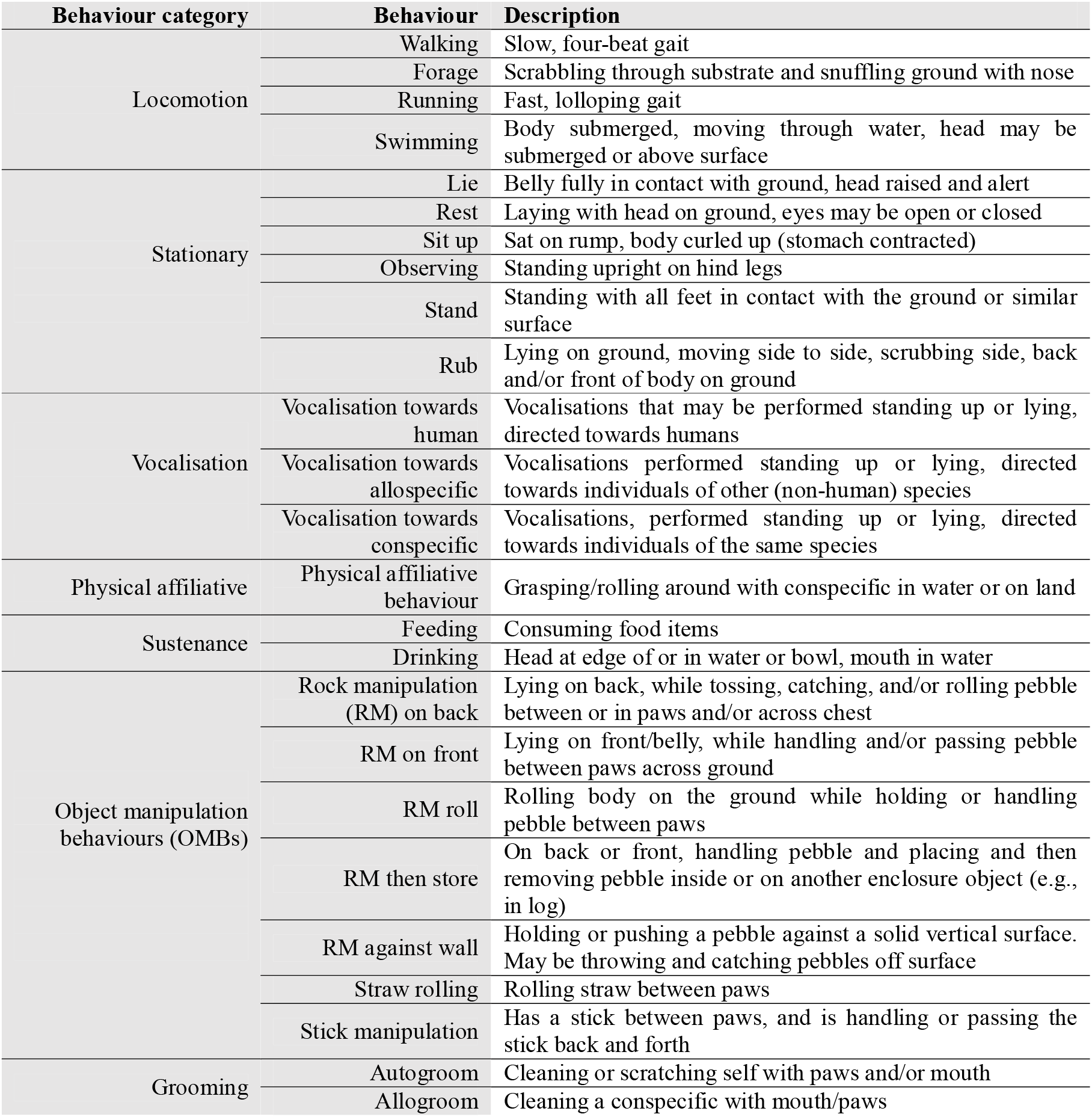
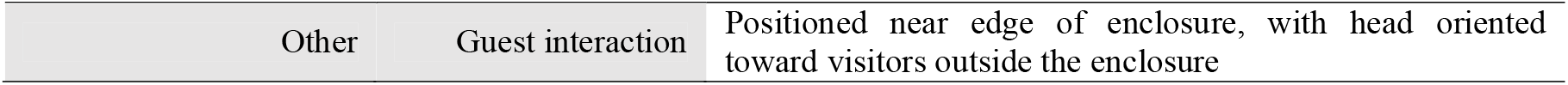
Ethogram to characterise object manipulation and broader behavioural repertoire observed in the Asian small-clawed otters (n=6) housed at the Lincolnshire Wildlife Park. Adapted from Allison et al. (2020).

#### 2.3.3 Coding

The ethogram was coded in BORIS (Behavioral Observation Research Interactive Software) for key press recording of behaviour. Continuous focal following was used to record the number (n) of bouts of behaviours and the duration of behaviours (s) per bout. A bout was defined as a single continuous display of a behaviour from start to finish; consecutive bouts were determined as distinct if separated by more than 2 seconds. Bouts may have spanned multiple consecutive video clips. In this study, it was not possible to accurately identify individual otters and therefore whilst data were recorded per animal in each video record, behaviours were not assigned to any particular individual (see Section 4.1 Limitations).

### 2.4 Weather and visitor density

Precipitation and air temperature data for the study period were obtained from the Met Office Weather Observation Website (https://wow.metoffice.gov.uk/). Data from the three nearest stations were collected and a distance-weighted average used to calculate mean daily temperature (stations: Wainfleet, Louth, Mareham) and total daily rainfall (stations: Leverton Highgate, Ulceby Cross, Gibraltar Burgh Sluice). Due to there being very little rainfall during the study period, only air temperature data were used in analyses. Online ticket sales were used as a proxy for total visitor numbers, as daily walk-in admittance data were unavailable.

### 2.5 Statistical analysis

As it was not possible to identify individual otters for the purposes of statistical analysis each bout was treated as independent to enable some degree of statistical analysis. However, results are not strictly independent, as the same otters likely appear across numerous bouts at unknown frequencies; this is acknowledged as pseudoreplication and thus results should be interpreted with this limitation in mind.

Kruskal-Wallis tests were used to assess differences in daily bouts (n) and duration (s) of OMBs across different cameras. Spearman’s corelation was used to assess the relationships between OMBs and visitor numbers (n), daily precipitation and mean daytime temperature (°C). A generalized linear model with a Tweedie distribution was used to assess the association of daily visitor numbers (n) and mean daytime air temperature (°C) on the total daily duration of OMBs.

Principal component analysis (PCA) was performed to explore the expression of and repertoire of behaviours expressed across the different cameras/locations. Analysis was performed at the ‘behaviour category’ level (see Table 1) and computed based on the total time spent performing behaviours in that category, by observation. PCA results were visualised using biplots, with observations categorized based on the camera/location from which they were obtained.

## 3 Results

In total, 2985 video clips were recorded. Of these, 1054 (36.4%) were from the Rock Cam, 697 (24.1%) from the Feed Cam, and 1144 (39.5%) from the Den Cam.

### 3.1 Descriptive summary

Otters were most frequently observed performing behaviours within the Locomotion and Stationary behavioural categories over the study period. The discrete behaviours most frequently observed were Walking (21.5%), followed by Lie (20.8%), and Feeding (12.2%) (Table 2).

**Table 2.**
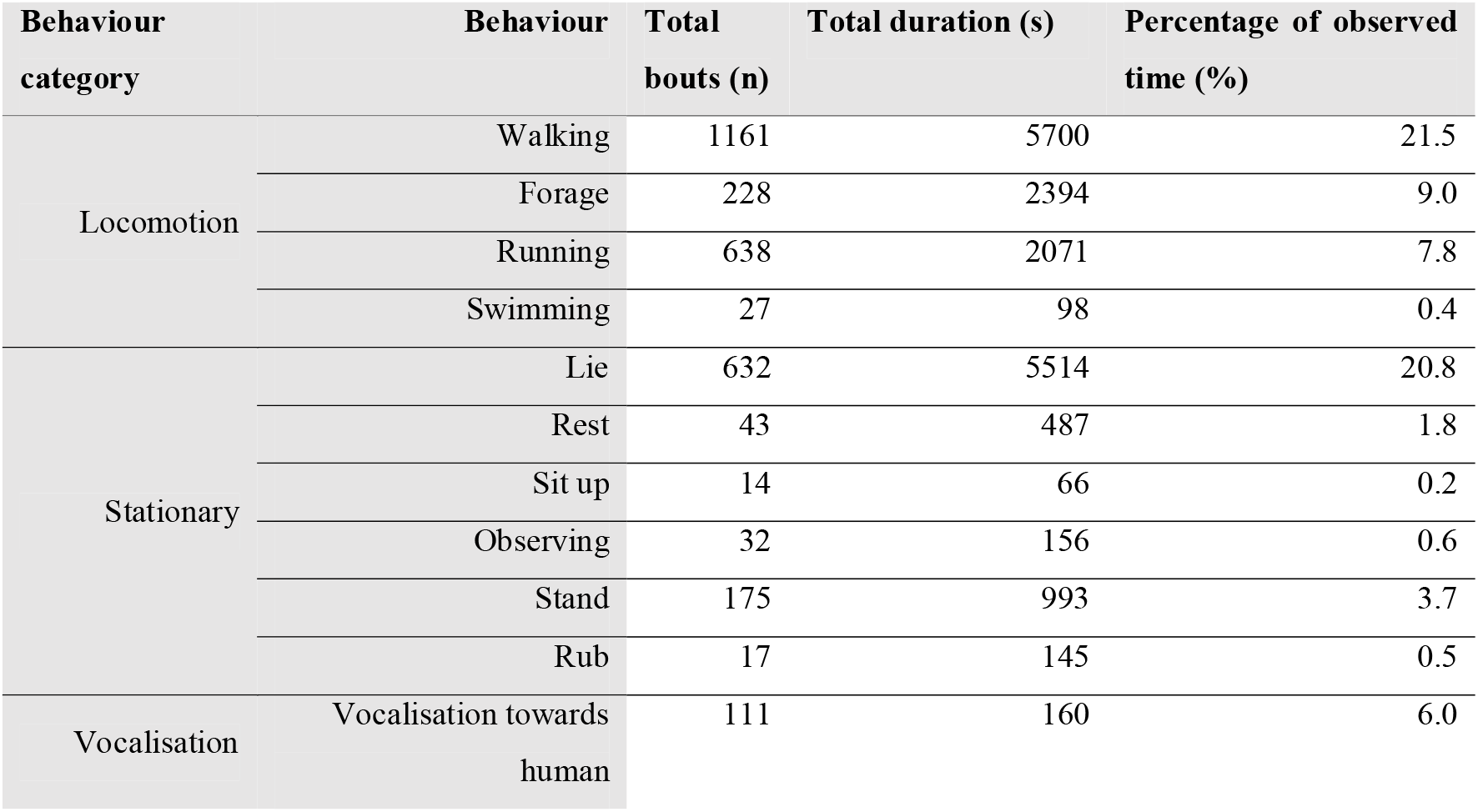
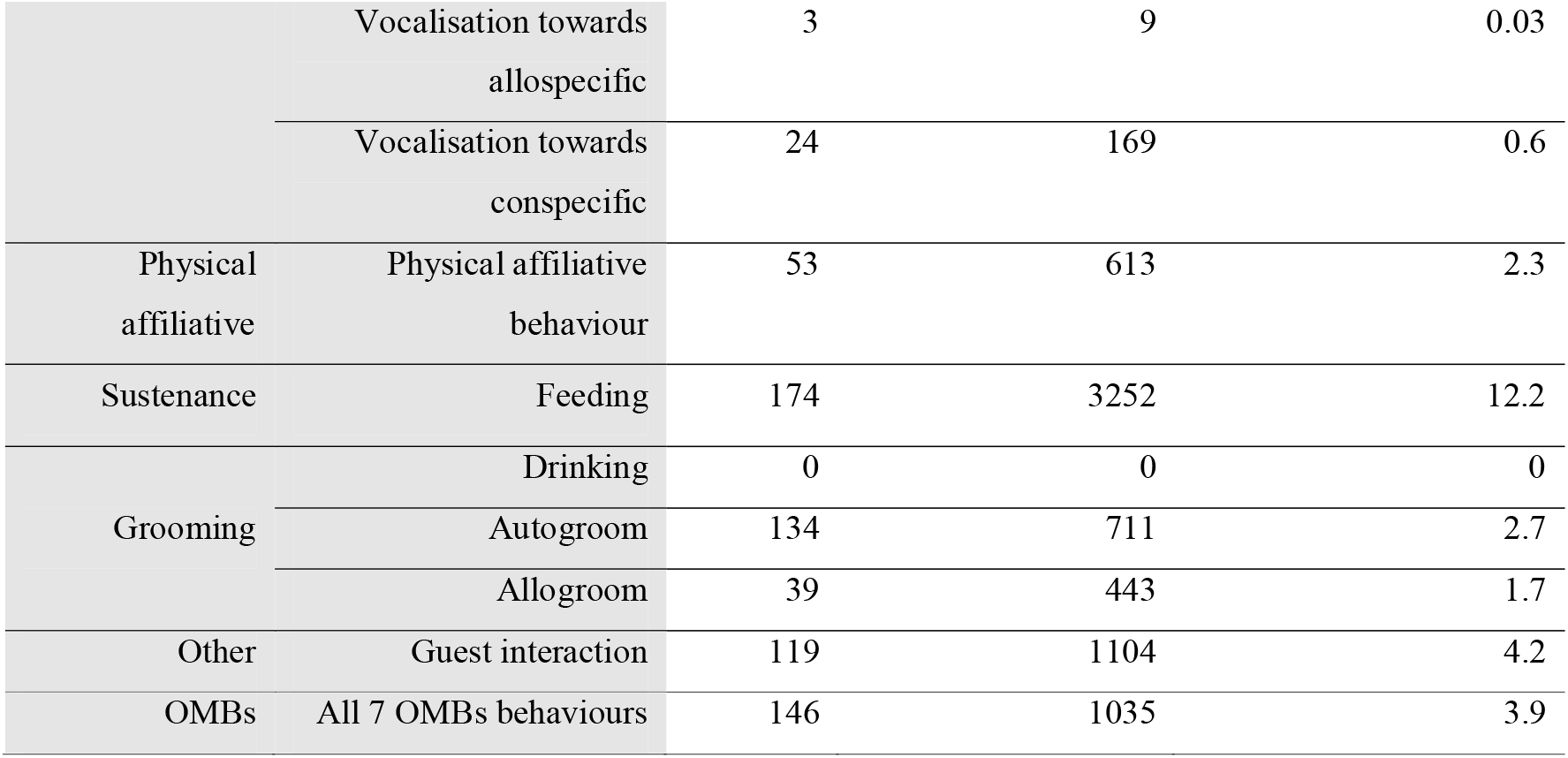
Total behaviour bouts (n), duration (s), and overall percent of study observation time (%) Asian small-clawed otters (n=6) were observed performing behaviours.

OMBs made up just 3.9% of the overall time budget recorded on the camera traps. Within the OMBs category, rock manipulation was most common, while straw and stick manipulation was relatively infrequent (Table 3). Rock manipulation was most often observed performed on the ground, with the two most common forms being rock manipulation (RM) on front (27.4% of time observed of all OMBs, followed by RM on back (22.8% of all OMBs) (Table 3).

**Table 3.** Total bouts (n) and duration (s) of object manipulation behaviours (OMBs) observed in Asian small-clawed otters (n=6) at Lincolnshire Wildlife Park during the study period.

| Behavioural<br>categories | Discrete behaviour | Total bouts (n) | Total duration (s) | Percentage of<br>time (%) |
| --- | --- | --- | --- | --- |
|  | Rock manipulation (RM) on back | 27 | 241 | 22.8 |
|  | RM on front | 44 | 290 | 27.4 |
|  | RM Roll | 17 | 145 | 13.7 |
| OMBs | RM then store | 9 | 62 | 5.9 |
|  | RM against wall | 18 | 154 | 14.6 |
|  | Straw rolling | 13 | 103 | 9.7 |
|  | Stick manipulation | 8 | 62 | 5.9 |

### 3.2 Factors associated with OMBs

The performance of OMBs varied across the three observed locations (Figure 3), both in bout number (□^2^ = 13.298, p = 0.001; mean daily number of bouts being 3.9 at Den Cam, compared to 0.4 at Rock Cam and 0.3 at Feed Cam) and in time spent observed performing OMBs (□^2^ = 13.047, p = 0.001; mean daily duration of 28.6s at Den Cam, compared to 3.9s at Rock cam and 2.2s at Feed Cam).

**Figure 3.**
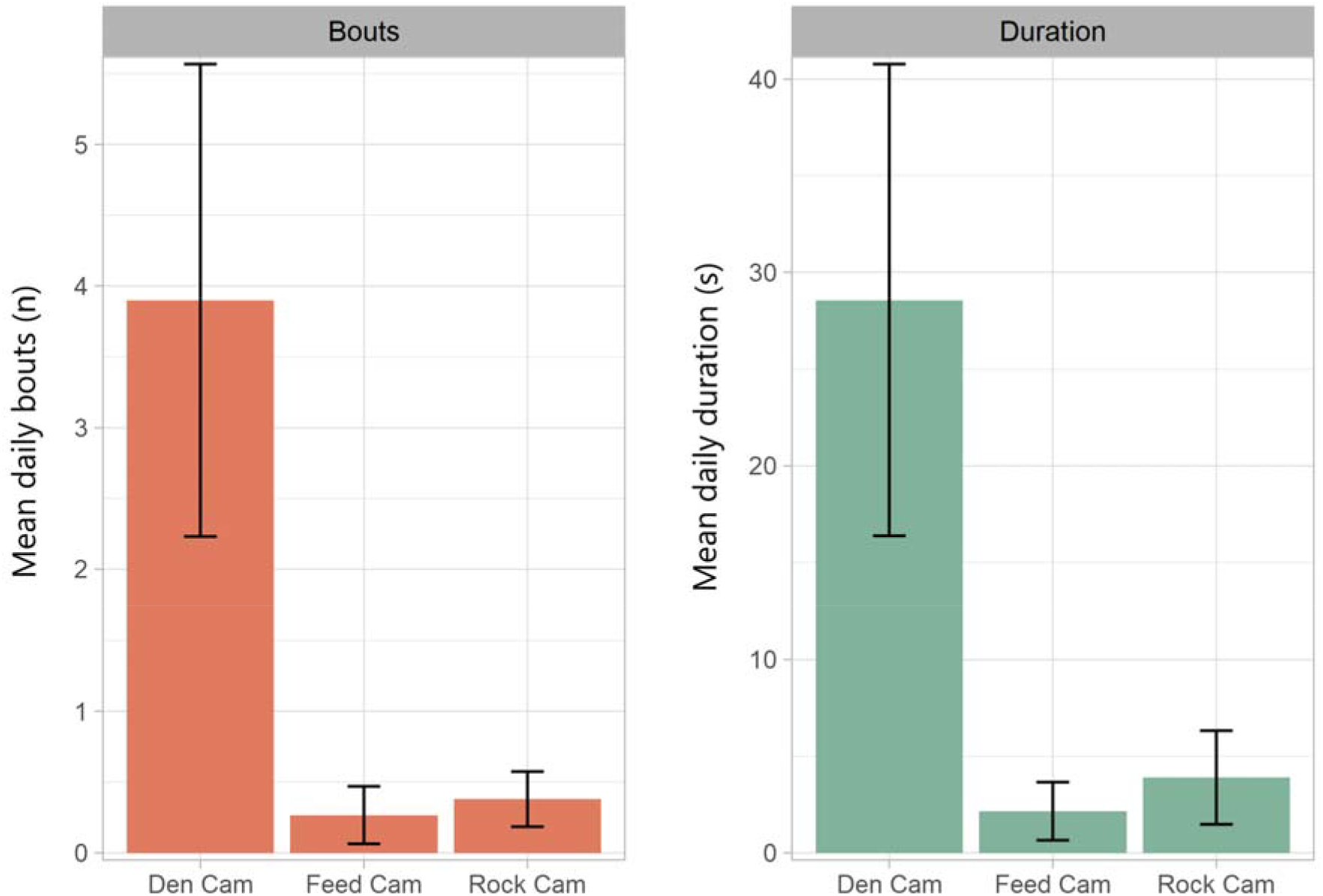
Mean daily bouts (n) and duration (s) of object manipulation behaviours (OMBs) across different camera sites within the Asian small-clawed otter (n=6) enclosure. Error bars represent standard error.

The general linear model showed a negative association of air temperature with OMBs (*t* = - 4.388, p < 0.001) but there was no significant association of visitor numbers with OMBs (*Z* = -0.811, p = 0.424).

### 3.3 Principal component analysis

For the PCA of categorised behaviours, the majority of variance was explained by PC1 (40.19%) and PC2 (23.36%) (Table 4, Figure 4). PC3 and PC4 explained far less variance, but specific factors, such as sustenance, were best characterised in these (Table 4, Figure 5).

**Table 4.** Loadings of factors / behaviour categories in principal component analysis of Asian small-clawed otter (n=6) behaviours.

|  | PC1 | PC2 | PC3 | PC4 |
| --- | --- | --- | --- | --- |
| Locomotion | 0.476 | -0.173 | -0.147 | -0.510 |
| Stationary | 0.469 | -0.095 | 0.002 | 0.107 |
| Vocalisation | 0.490 | -0.353 | 0.114 | 0.175 |
| Phys. Affiliative | 0.453 | -0.274 | 0.185 | 0.166 |
| Sustenance | 0.192 | 0.245 | -0.879 | 0.254 |
| Grooming | 0.120 | 0.585 | 0.305 | 0.663 |
| OMBs | 0.240 | 0.599 | 0.257 | -0.408 |
| Variance explained | 40.19% | 23.36% | 13.36% | 9.08% |

**Figure 4.**
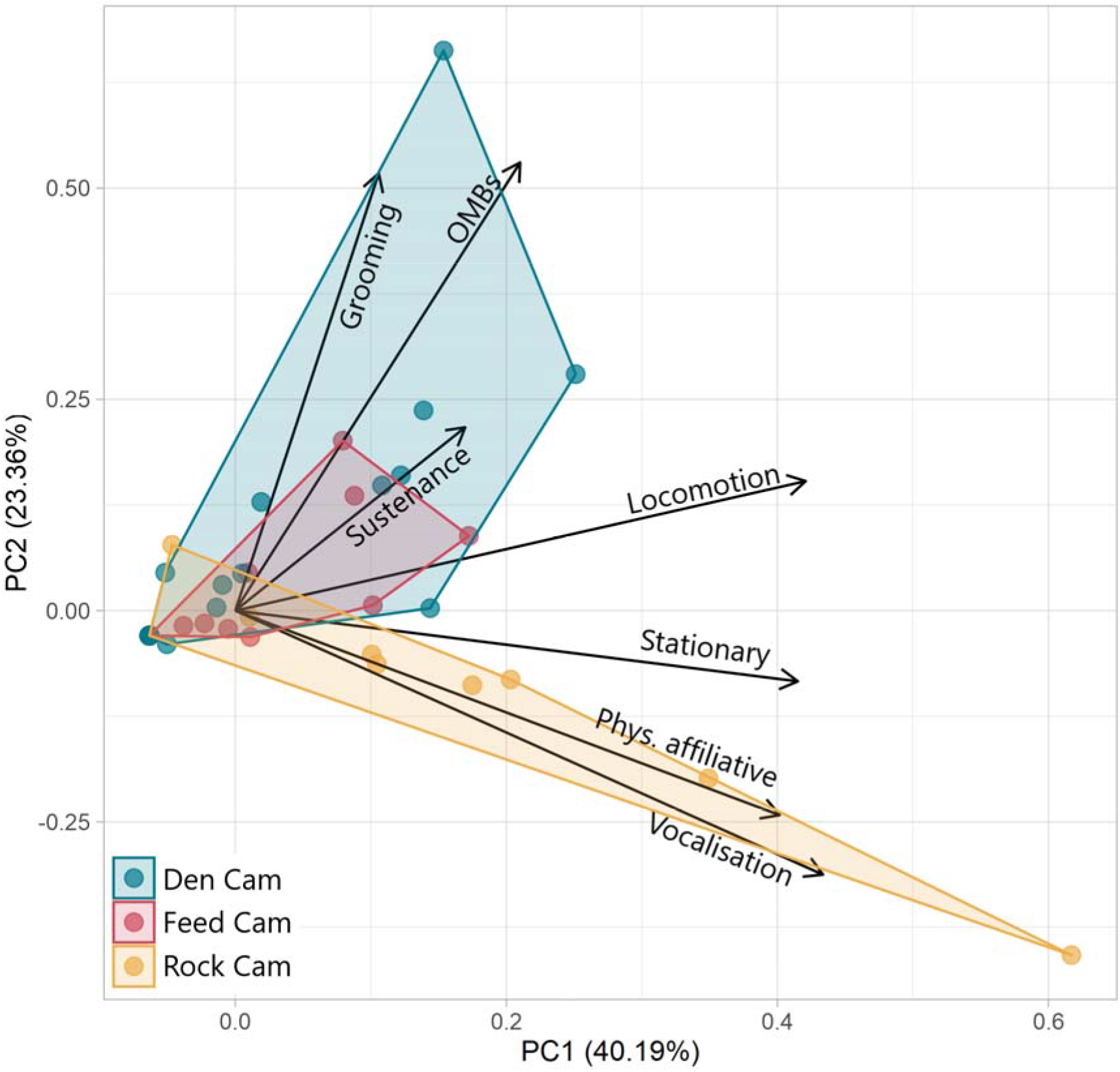
Biplot of PC1 and PC2 from principal component analysis (PCA) of categorised behaviours in Asian small-clawed otters (n=6), grouped by camera / enclosure location.

**Figure 5.**
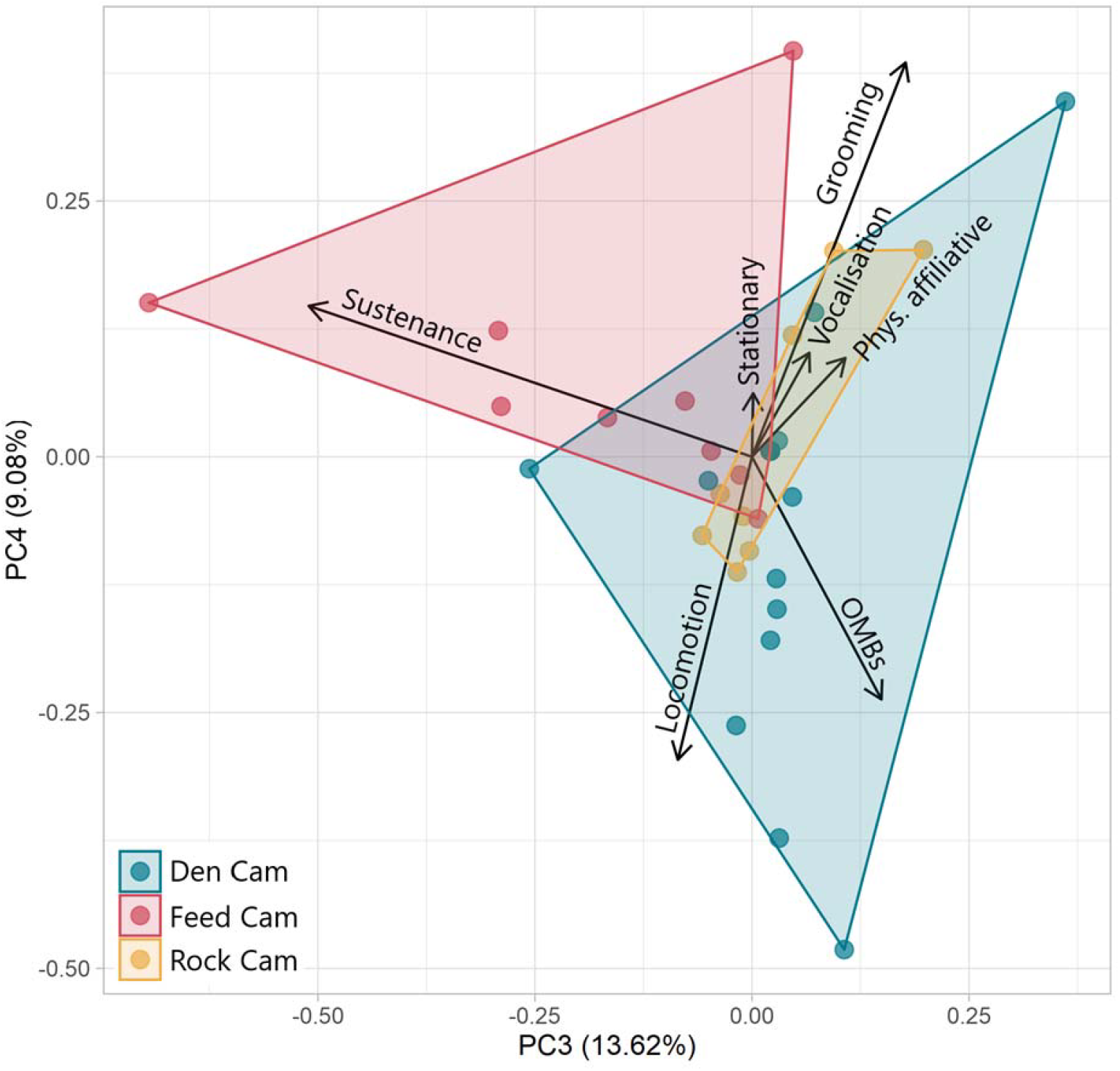
Biplot of PC3 and PC4 from principal component analysis (PCA) of categorised behaviours in Asian small-clawed otters (n=6), grouped by camera / enclosure location.

**PC1** was characterised by moderate loadings for locomotion (0.476), stationary (0.469), vocalisation (0.490), and physical affiliative (0.454). The greatest variation in the expression of these factors was at the Rock Cam.

**PC2** had especially strong loadings for OMBs (0.599) and autogrooming (0.585), both of which were primarily associated with the Den Cam.

**PC3** had a single predominant loading of ‘sustenance’ (-0.879), a factor which only featured lightly in PC1 (0.192) and PC2 (0.245). This was strongly associated with the Feed Cam and there was a clear gradient of observations on that camera along this factor.

**PC4** had moderately strong positive loading for grooming (0.663), but negative loadings for locomotion (-0.510), and OMBs (-0.408), which contrasts with PC2, albeit explaining a much lower % of variation that PC2.

## 4 Discussion

Object manipulation behaviours have been widely reported in wild and captive animal populations (Breland and Breland 1961; Huffman and Quiatt 1986). In wild populations, it is generally assumed the behaviours have some adaptive value, for example in exploitation of environmental resources such as food acquisition or in the development and refinement of skills. An example of the development of such behaviours are the Japanese macaques (Huffman and Quiatt 1986; Leca et al. 2007; Nahallage and Huffman 2007), who have been documented to have developed several food processing skills such as potato washing (Kawai 1965) when their diet was supplemented by additional food resources. These behaviours demonstrated both functional benefits in terms of food exploitation, as well as evidence of learning through social transmission within the population (de Waal and Bonnie 2009). It is noteworthy that these populations had also been observed to perform object manipulations with non-food items such as stones (Leca et al. 2007), which were suggested to be part of this skill development.

Whilst adaptive value can be ascribed to the expression of object manipulation behaviours in captive populations, it has also been suggested that these behaviours may be related to inappropriate or poor environmental conditions, where animals may not be able to fully express patterns of behaviour found in the wild (Wood-Gush and Vestergaard 1989). In this context, the activities may represent a form of displacement behaviour, due to motivational conflicts and/or redirected behaviour where suitable resources are not available for the conventional expression of motivated activities. In extreme circumstances they may develop into repetitive, stereotypic responses that may indicate a welfare concern in captivity (Troisi 2002; Mason and Latham 2004; Rodenburg et al. 2004). For example, dry sows in outdoor production systems commonly develop a stone-chewing behaviour (Lawrence and Terlouw 1993; Horrell and A’Ness 2003), which is analogous to stereotypic patterns of bar-biting observed in stall-housed dry sows and gilts (Dantzer and Mormede 1983; Appleby and Lawrence 1987). In both cases the behaviours have been associated with prolonged food deprivation and are consistent with redirection of feeding and foraging activity to non-nutritive objects by hungry animals (Lawrence and Terlouw 1993).

In previous work involving captive otters, Allison et al. (2020) concluded that the activities may be best described as an indication of hunger, with otters misdirecting feeding and foraging behaviour to non-nutritive resources. This conclusion was primarily related to the timing of these behaviours in their study populations, with fewer observations of rock-juggling behaviour in the two hours following feeding times when otters were assumed to be satiated and compared with periods more than two hours after feeding times when otters were assumed to be hungrier. Although the authors suggested this temporal distribution of the behaviours may be evidence of hunger as the underlying motivation, they did not conclude that hunger was the only contributing factor and recommended further work to better understand underlying causes. In our study, however, we found no evidence of a direct association between feeding behaviour and object manipulation: rather, object manipulation was not commonly observed around feeding time but was commonly observed in an area of the exhibit where the otters were not normally fed. This suggests a different causal basis in this population at least, with the timing and location of the behaviour more consistent with maintenance or play behaviours.

In our study, OMBs were predominantly observed around the otters’ den. This area was partially worn bare of grass, suggesting a high degree of activity in this area, which keepers also reported. Grooming behaviour was also frequently observed here, which tends to be expressed during periods of lower arousal. Although grooming and other self-directed behaviours have been described as forms of displacement activity when performed in an excessive manner or out of context (Delius 1967), they are normally a form of self-care, performed during periods of lower arousal, such as in the boundaries between environmental interaction and resting behaviour (Cohen and Price 1979). In our study, the grooming behaviour was consistent with self-care during periods of relaxation, being performed away from main public view in proximity to the den and not expressed alongside activities that may represent higher levels of arousal such as locomotion, feeding and vocalisations. If OMBs had been driven by hunger in our population, we would have expected a greater expression of these behaviours at the feeding area at the front of the enclosure.

It is possible that the feeding regime at LWP may mean that the otters do not experience hunger to the extent necessary for it to influence behaviour. In addition, the zoo did not advertise its feeding times for the otters, which could otherwise result in the accumulation of visitors around the enclosure vicinity and resultant development of apparently anticipatory activities such as pacing or vocalisations in captive carnivores (Mason 1991). Furthermore, the enclosure at LWP is relatively large, diverse, and enriched, which may have the effect of reducing the experience of hunger and potential re-direction of feeding/foraging behaviour to non-nutritive objects (Mason et al. 2007; Grimm and Sauter 2020). In short, the feeding regime for our study population may not have resulted in high motivation to persevere or misdirect feeding behaviour; where otters may have been motivated to forage/seek additional food, they had a large and variable environment in which this foraging behaviour could be expressed.

An alternative explanation for OMBs could lay in the possibility of their classification as displacement behaviours, which happens during periods of social tension and can cause negative affective states such as stress, anxiety, and frustration (Troisi 2002). It is noteworthy that other behaviours commonly seen in captive carnivores, such as pacing or excessive vocalisations (Mason and Latham 2004; Mason et al. 2007), were not observed in this population; while singular measures of welfare should be interpreted cautiously, the lack of these behaviours here suggests that stress may have been largely mitigated in the otters’ large, varied enclosure.

Though multivariate analysis showed no significant association of visitor numbers with OMBs, we would like to highlight that a univariate (Spearman’s) correlation did suggest a negative association (*r* = -0.421, *p* = 0.021). We highlight this as something worth future consideration, not least given the limitations of our analyses. If there was such an effect, this may inform a better explanation as to why otters were most often observed performing OMBs near their den compared to the other areas: the area around the den was the most occluded from the view of visitors. This would be consistent with the findings of Rossi et al. (2020), who found that otters’ instances of ‘locomotion’ and ‘play’ increased during closed periods when compared to periods of time when the zoo was open. This explanation also aligns with the wider implication that absence of play may be an indicator of poor welfare in captive animals (Held and Špinka 2011). If these behaviours had been seen in a wild population of otters, then it would be conventional to consider the behaviours to be part of an animal’s normal behavioural repertoire and hold potential adaptive value such as acquisition and practise of manipulative skills. As the evidence in our study was not consistent with redirection of a prevented behaviour or the expression of a displacement activity due to motivational conflict, it is worth considering if object manipulation could be better explained as the expression of normal behaviour patterns directed at available resources--in other words, a form of play behaviour. Here then, the den area may be one in which otters putatively feel safe and relaxed, which would also explain the relatively higher incidence of grooming in that location. In other animals, play and grooming have been associated with states of relaxation and areas of safety (Spruijt et al. 1992; Feh and de Mazières 1993; Held and Špinka 2011). Supportive of this conclusion is that our observations appear to meet Burghardt’s (2005) five criteria for play:

1. **Non-functional**: Behaviours such as straw rolling appeared to not contribute to meeting any basic needs (i.e. feeding, mating) of the otters. They often abandoned the substrate immediately after use.
2. **Spontaneous and pleasurable:** We did not observe OMBs performed in response to any apparent stimuli, so could consider these to have occurred spontaneously. The behaviours appeared to be freely performed, indicating that they may be pleasurable. Certainly, when observing the otters expressing some of these behaviours (e.g. rubbing while rock manipulating), it does appear that performance of these appears to *feel good*.
3. **Incomplete or exaggerated:** The ranges of motion were more extreme and pronounced than for other handling activities, such as foraging. For example, rocks would often be fumbled, dropped, and re-handled multiple times during any event. Otters would roll and writhe whilst performing OMBs.
4. **Repetition:** The behaviours were repeated continuously and repeatedly within any individual event, but also repeated over the study period.
5. **Good physiological state:** The otters were well-fed, receiving regular meals, and in overall good health. The nature of OMBs is that otters were often laying on their backs, making movements and sounds that would alert predators. It can be surmised that they would be unlikely to perform such behaviours unless they felt safe and relaxed.

### 4.1 Study considerations and limitations

Several study artifacts impact the interpretation of our findings. First, this study was conducted during the summer. We found that OMBs decreased as air temperature rose; this is inconsistent with the increase in most behaviours (though not stereotypies, which decreased) observed during warmer months in another group of Asian small-clawed otters housed indoors in the UK (Cuculescu et al. 2017); it is possible the otters in our study experienced temperature shifts more acutely as a result of being housed outdoors, but we also note that as data were only collected in the summer months, we cannot explain how colder temperatures may impact expression of these behaviours. Second, online ticket sales were used as a proxy to approximate visitor density; while others (Mooney et al. 2020) have used similar approaches, more sophisticated measures of visitor density and behaviour around the enclosure would better elucidate the impact of visitors on OMBS in otter populations.

Finally, the primary limitation of this study is potential for pseudoreplication in analyses. Individual bouts were considered as independent observations. However, this is not strictly true, as each otter will have been present in an unknown number of observations. Furthermore, bouts may influence each other, for example if an otter has groomed earlier in the day, it may be less inclined to groom in the near future. We stress the importance of this to readers when interpreting results. Issues of pseudoreplication are quite common in zoo studies, which often observe one, or few, populations (Waller et al., 2013; Pastorino et al., 2021; Bandoli et al., 2023). Identification of individual otters and the inclusion of additional otter troops would facilitate a more comprehensive and representative analysis.

### 4.2 Conclusions

Our findings suggest that in this population of otters, object manipulation did not appear to be a response to environmental stressors associated with captivity, or to frustration of important behaviours such as feeding and foraging. We tentatively suggest that the behaviours observed amongst the otters in our study population may be a form of play, involved in the development and practise of manipulative skills. An innovative approach used in this study has been the application of multi-variate analysis, not only for data reductions but to investigate associations between different behaviours to better understand the motivations underlying development of novel behaviour patterns. This has the potential to compliment other approaches to investigate relationships between different behaviours using controlled, experimental manipulations of animal management to further test underlying causes of apparently abnormal responses to captivity. In this context, systematic investigation of the role of feeding regimes and the impact of external factors (e.g., visitors, weather) on the incidence of behaviours that may be associated with positive and negative affective state would be a valuable direction for future research.

## Supplementary files for this manuscript are available at

https://osf.io/gup9j/overview?view_only=7bc5525491e24b669f620f7ccfd10cb5

## Conflict of Interest

The authors declare no conflicts of interest.

## Notes

https://osf.io/gup9j/overview?view_only=7bc5525491e24b669f620f7ccfd10cb5

### Competing Interest Statement

The authors have declared no competing interest.

